# Glycine targets NINJ1-mediated plasma membrane rupture to provide cytoprotection

**DOI:** 10.1101/2021.12.12.471765

**Authors:** Jazlyn P. Borges, Ragnhild S. R. Sætra, Allen Volchuk, Marit Bugge, Bridget Kilburn, Neil M. Goldenberg, Trude H. Flo, Benjamin E. Steinberg

**Author notes:** Equally contributing first authors (Jazlyn P. Borges, Ragnhild S. R. Sætra). Corresponding author: Benjamin E. Steinberg, Peter Gilgan Centre for Research and Learning, Hospital for Sick Children, 686 Bay Street, rm 06-9707, Toronto, Ontario, Canada, M5G 0A4, E.

## Abstract

First recognized 35 years ago, glycine is known to protect cells against plasma membrane rupture from diverse types of tissue injury. This robust and widely observed effect has been speculated to target a late downstream process common to multiple modes of tissue injury. The molecular target of glycine cytoprotection, however, remains entirely elusive. We hypothesized that glycine targets ninjurin-1 (NINJ1), a newly identified executioner of plasma membrane rupture in pyroptosis, necrosis, and post-apoptotic cell death. NINJ1 is thought to cluster within the plasma membrane to cause cell rupture. Here, we first demonstrate that NINJ1 knockout functionally and morphologically phenocopies glycine cytoprotection in mouse and human macrophages stimulated to undergo lytic cell death. Next, we show that glycine treatment prevents NINJ1 clustering thereby preserving cellular integrity. By identifying NINJ1 as a glycine target, our data help resolve a long-standing mechanism of glycine cytoprotection. This new understanding will inform the development of cell preservation strategies to counter pathologic lytic cell death pathways.

**Graphical abstract:** 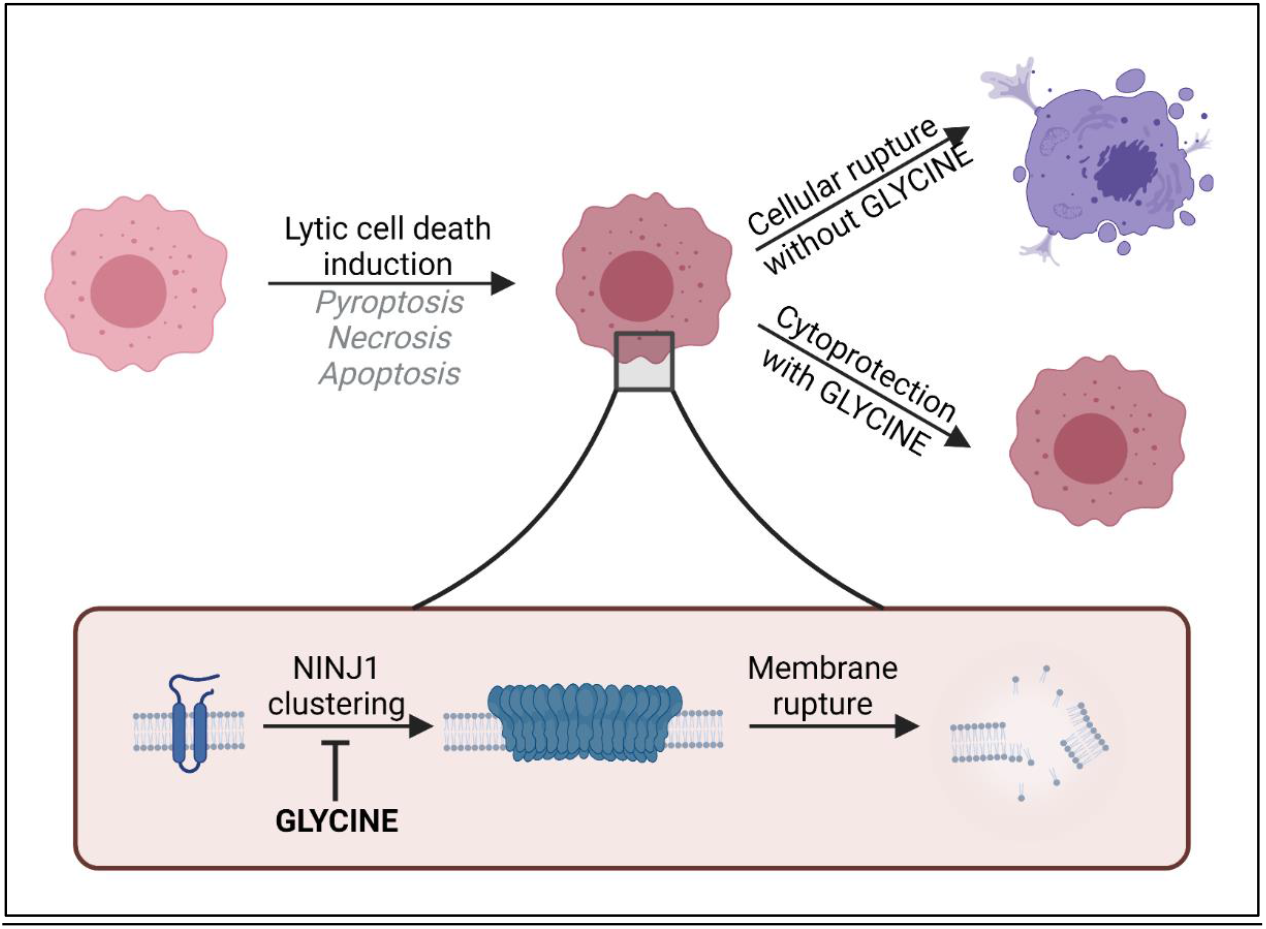

**Summary:** Glycine is known to protect cells against plasma membrane rupture from diverse types of tissue injury by an unknown mechanism. The authors demonstrate that NINJ1, a newly identified executioner of plasma membrane rupture across lytic cell death pathways, is a glycine target and resolve a longstanding mechanism of glycine cytoprotection.

## Introduction

The amino acid glycine has long been known to protect cells against plasma membrane rupture induced by a diverse set of injurious stimuli. This cytoprotective effect was first described in 1987 where glycine was found to protect renal tubular cells against hypoxic injury (Weinberg et al., 1987). Since, similar observations have been replicated across immune, parenchymal, endothelial and neuronal cell types in multiple injury models [reviewed in (Weinberg et al., 2016)]. These include the lytic cell death pathways pyroptosis (Loomis et al., 2019; Volchuk et al., 2020) and necrosis (Estacion et al., 2003).

While the robust cytoprotective effects of glycine are widespread, the molecular target and mechanism by which glycine protects cells and tissues remain elusive. Many cellular processes have been investigated but ultimately ruled out as underlying this phenomenon (Weinberg et al., 2016). These include glycine or ATP metabolic pathways (Weinberg et al., 1991), intracellular calcium or pH regulation (Weinberg et al., 1994), cytoskeleton stabilization (Chen et al., 1997), and chloride conductance (Loomis et al., 2019). It has been speculated that glycine targets a membrane receptor but is unlikely to involve canonical glycine receptors (Weinberg et al., 2016; Loomis et al., 2019). Lastly, as a cytoprotective agent, glycine has oftentimes been described as an osmoprotectant, but its small size generally allows free passage through the large membrane conduits involved in lytic cell death pathways, such as the gasdermin D pores formed during pyroptosis (Xia et al., 2021), thereby negating a protective osmotic effect. A mechanistic knowledge of how glycine preserves cellular integrity, however, would inform the development of cell and tissue preservation strategies with clear clinical implications, such as for ischemia-reperfusion injuries where multiple lytic cell death pathways are activated to cause tissue damage and morbidity (Del Re et al., 2019).

Given how widely observed the phenomenon is, it stands to reason that glycine targets a late downstream process common to multiple tissue injury models. Notably, the transmembrane protein ninjurin-1 (NINJ1) was recently identified as the common executioner of plasma membrane rupture in pyroptosis, necrosis, and post-apoptosis lysis (Kayagaki et al., 2021). Post-apoptosis lysis refers to the cell membrane rupture that occurs when phagocytes are unable to scavenge apoptotic cells that have otherwise completed the apoptotic program (Silva, 2010). NINJ1 is a 16-kilodalton protein with two transmembrane regions that resides in the plasma membrane and is reported to function as a cell adhesion molecule (Araki et al., 1997; Araki and Milbrandt, 1996). It was first identified for its role in nerve injury (Araki and Milbrandt, 1996) and has since been implicated in multiple inflammatory conditions, such as atherosclerosis, ischemic brain injury and tumorigenesis (Jeon et al., 2020; Kim et al., 2020; Yang et al., 2017). Its involvement in lytic cell death pathways involves its clustering within the plasma membrane, which results in cellular disruption by an unknown mechanism (Kayagaki et al., 2021).

Here, we hypothesized that glycine targets NINJ1 to mediate its cytoprotective effect. We first demonstrate that NINJ1 knockout or silencing functionally and morphologically phenocopies glycine cytoprotection in mouse and human primary macrophages stimulated to undergo various forms of lytic cell death. Next, we show that glycine treatment prevents NINJ1 clustering within the plasma membrane, thereby preserving its integrity. By identifying NINJ1 as a glycine target, our data help resolve a long-standing mechanism of glycine cytoprotection.

## Results and Discussion

### NINJ1 deficiency functionally and morphologically phenocopies glycine cytoprotection

The published literature suggests a potential association between NINJ1 and glycine given that NINJ1 knockout and glycine treatment protect cells against a common set of lytic cell death pathways. Moreover, in pyroptosis, both allow for IL-1β secretion through the gasdermin D pore while preventing final membrane lysis (Volchuk et al., 2020; Kayagaki et al., 2021). We first sought to extend these associations by further characterizing the functional and morphological similarities between NINJ1 knockout and glycine cytoprotection in pyroptosis, necrosis and post-apoptosis lysis.

Using a CRISPR-Cas9 system, we generated a NINJ1 knockout cell line in immortalized bone marrow-derived macrophages (iBMDM) (**Figure 1A**). Pyroptosis was induced in LPS-primed wildtype and NINJ1 knockout iBMDM with nigericin (20 μM, 2 hrs) in the absence or presence of 5 mM glycine. We performed a colorimetric assay to test for lactate dehydrogenase (LDH) release, a common marker of cell rupture. Both NINJ1 knockout and glycine treatment in wildtype iBMDM protected against cytotoxicity (**Figure 1B**). Similarly, cell membrane integrity was preserved following apoptosis (**Figure 1C**) and necrosis (**Figure 1D**) induced with venetoclax (25 μM, 16 h) and pneumolysin (0.5ug/ml, 15 min), respectively. Importantly, in NINJ1 knockout macrophages, treatment with glycine during lytic cell death did not confer any additional protection against cell lysis (**Figure 1B-D**).

**Figure 1.**
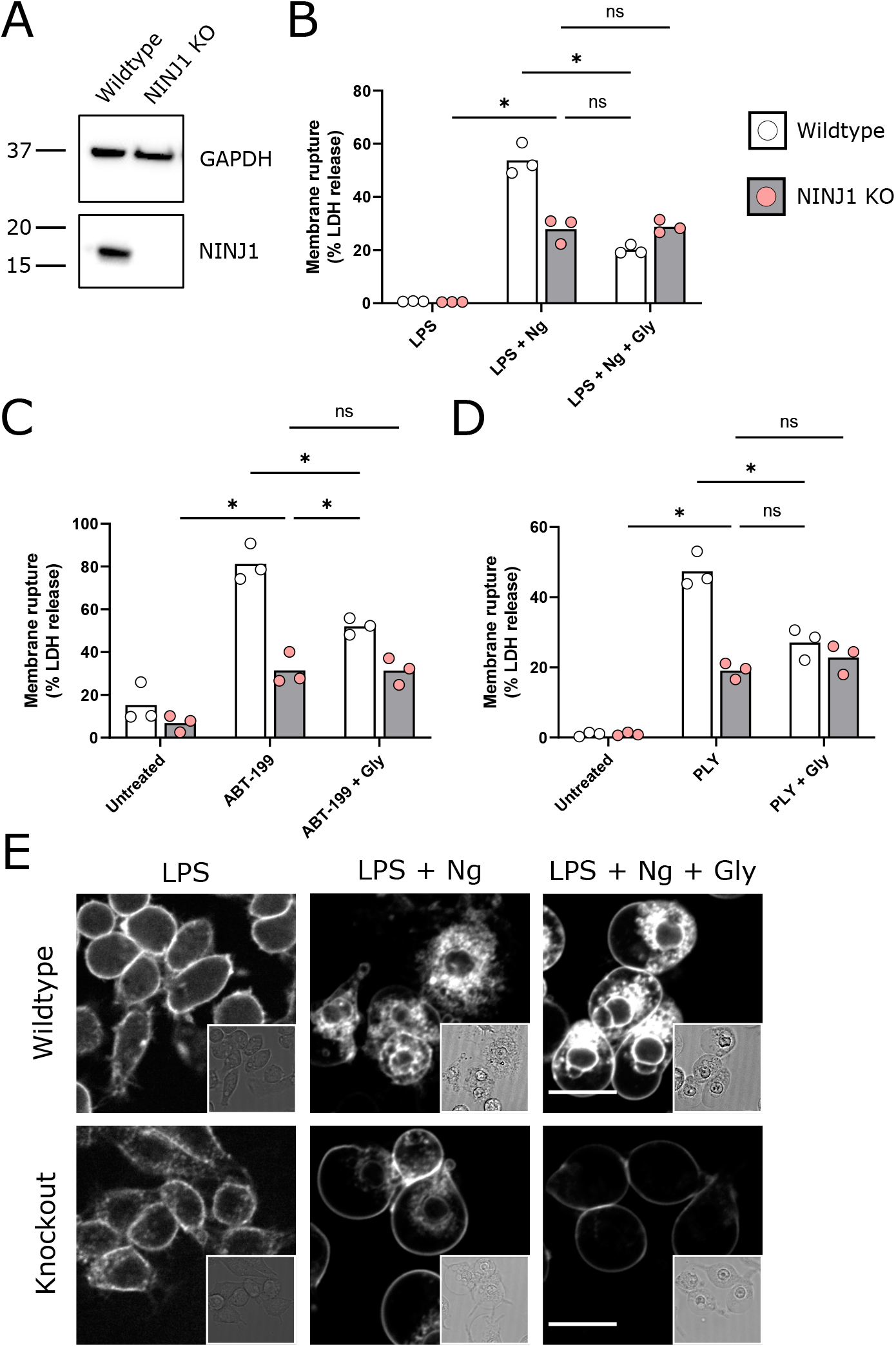
NINJ1 knockout functionally and morphologically phenocopies glycine cytoprotection. (**A**) Immunoblot analysis demonstrating NINJ1 knockout in iBMDM macrophages. GAPDH is presented as a loading control. (**B-D**) Wildtype and NINJ1 knockout iBMDM were induced to undergo pyroptosis (LPS + nigericin), secondary necrosis (venetoclax) or necrosis (pneumolysin) with or without 5mM glycine treatment. Cytotoxicity was evaluated by measuring the levels of LDH in the supernatant. Cytotoxicity decreased in glycine-treated wildtype cells comparably to NINJ1 knockout across each of (**B**) pyroptosis, (**C**) secondary necrosis, and (**D**) necrosis. Glycine treatment of NINJ1 KO cells provided no additional protection to knockout cells treated without glycine. Data are expressed as supernatant LDH as a % of total LDH from lysates and supernatants, from *n* = 3 independent experiments for each cell death type. Individual data points are shown along with their mean. * *P* < 0.05 by ANOVA with Tukey’s multiple comparison correction. (**E**) LPS-primed wildtype iBMDM induced to undergo pyroptosis in the presence of glycine demonstrate similar plasma membrane ballooning to NINJ1 knockout iBMDM induced to undergo pyroptosis. Membrane ballooning is shown in live cells labelled with the plasma membrane dye FM4-43. The corresponding brightfield image is shown in the inset. Scale bar 15 μm.

When visualizing glycine-treated pyroptotic macrophages, we observed prominent plasma membrane ballooning (**Figure 1E**). This impressive morphology was comparable to that of LPS-primed NINJ1 knockout macrophages stimulated to undergo pyroptosis with nigericin, which we (**Figure 1E**) and others (Kayagaki et al., 2021) have observed. Together, these functional and morphologic similarities buttress the association between glycine cytoprotection and NINJ1-mediated plasma membrane rupture.

To further corroborate the above findings, we conducted pyroptosis assays in RAW mouse macrophages. NINJ1 was knocked out using a CRISPR-Cas9 system and compared with the parental line (**Supplemental Figure 1A**). Pyroptosis stimulation in the NINJ1 mouse knockout functionally (**Supplemental Figure 1B**) and morphologically (**Supplemental Figure 1C**) phenocopied glycine treatment as seen in the iBMDM.

Human NINJ1 is 90% homologous to the mouse protein and was shown to be toxic when overexpressed in HEK293T cells (Kayagaki et al., 2021), but it is not known if endogenous NINJ1 is activated and mediates plasma membrane rupture in human cells undergoing programmed cell death. We knocked down NINJ1 in human primary monocyte-derived macrophages (hMDMs) using siRNA specific to NINJ1 (siNINJ1) and compared to non-targeting control siRNA (siCtrl). Knockdown efficiency was around 70% (**Supplemental Figure 2A)**. Both NINJ1 knockdown and glycine independently and completely prevented pyroptosis (LDH release) induced by nigericin (20 μM, 2 hrs) in LPS-primed hMDMs, and no additional effect was seen with combined treatment (**Figure 2A**). Similar to mouse macrophages, both siNINJ1 and glycine sustained the ballooned morphology of pyroptotic hMDMs (**Figure 2B**). Together, these data using macrophage types from both mouse and human demonstrate that genetic deletion or silencing of NINJ1 functionally and morphologically parallels glycine-mediated cytoprotection.

**Figure 2.**
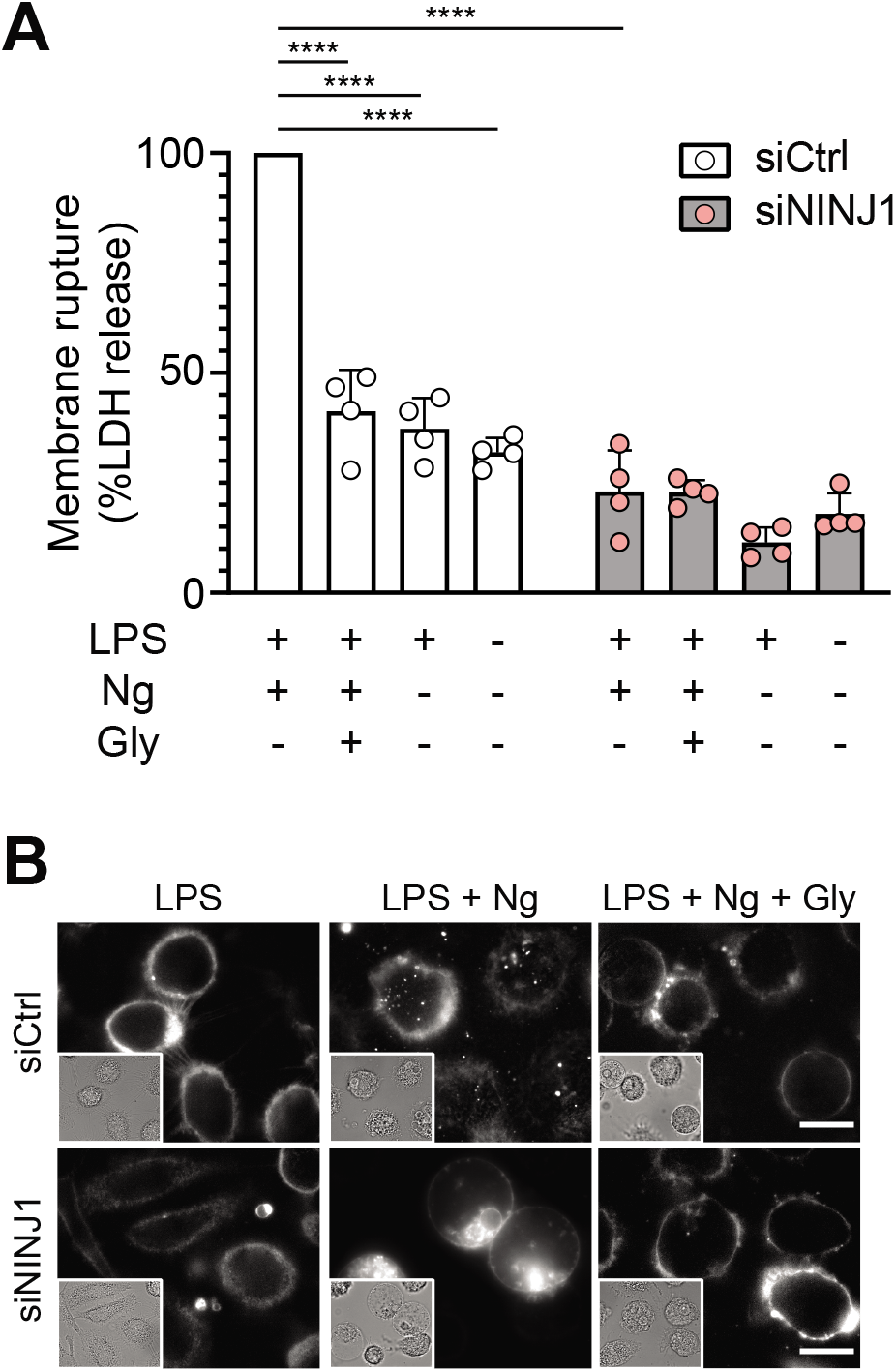
NINJ1 silencing in primary human monocyte-derived macrophages functionally and morphologically phenocopies glycine cytoprotection. Primary human monocyte-derived macrophages (hMDMs) were treated with siRNA targeting NINJ1 (siNINJ1) or non-targeted control (siCtrl). (**A**) Wildtype and NINJ1-silenced hMDMs were LPS-primed and induced to undergo pyroptosis (nigericin) with or without 50 mM glycine. LDH release in the supernatants was measured to assess cytotoxicity. Cytotoxicity was reduced in glycine-treated wildtype cells, as well as in NINJ1-silenced cells. Glycine treatment did not yield additional protection from pyroptotic cell death in NINJ1-silenced cells. Data are expressed as % of LDH in supernatant from siCtrl-treated cells stimulated to undergo pyroptosis from *n* = 4 independent donors. Data points from each donor is shown along with their mean and standard deviation. **** *P* < 0.0001 by 2-way ANOVA with Tukey’s multiple comparison correction. (**B**) Nigericin- and glycine-treated LPS-primed wildtype hMDMs show similar ballooning morphology as nigericin-treated LPS-primed NINJ1-silenced hMDMs. Membrane ballooning is shown in live cells labelled with the plasma membrane dye FM1-43. The corresponding brightfield image is shown in the inset. Scale bar 20 μm.

### Glycine targets NINJ1 clustering to prevent membrane rupture

We therefore next posited that glycine targets NINJ1 as part of its cytoprotective effect. The working model of NINJ1-mediated membrane rupture involves its clustering within the plasma membrane (Kayagaki et al., 2021). Point mutants (e.g. K45Q and A59P) that prevent membrane rupture also fail to cluster upon induction of pyroptosis (Kayagaki et al., 2021), further reinforcing a connection between the clustering process and the capacity of NINJ1 to rupture membranes. To establish whether glycine directly interferes with NINJ1-mediated cell death, we next evaluated the effect of glycine treatment on NINJ1 clustering during cell lysis in pyroptosis, post-apoptosis, and necrosis.

Using a biochemical native-PAGE approach that maintains native protein interactions, endogenous NINJ1 of otherwise untreated primary mouse bone marrow-derived macrophages (BMDM) migrates at approximately 40 kDa, potentially indicative of NINJ1 dimers or trimers in unstimulated cells (**Figure 3A**) and consistent with its known ability to form homotypic interactions (Bae et al., 2017). In response to pyroptosis (nigericin in LPS-primed BMDM), necrosis (pneumolysin), and apoptosis (venetoclax) induction in primary BMDM, the endogenous NINJ1 signal shifts to a high molecular weight aggregate, suggestive of a clustering process. Importantly, this shift is completely abrogated by glycine treatment (**Figure 3A**). Using this same native-PAGE approach, glycine treatment of primary hMDMs (**Figure 4A**) and mouse RAW-ASC cells (**Supplemental Figure 1D**) undergoing pyroptosis prevented the shift of NINJ1 to a high molecular weight band. Finally, glycine inhibited NINJ1 oligomerization and pyroptotic lysis (as measured by LDH release) in LPS-primed and nigericin-treated human macrophages derived from induced pluripotent stem cells (iPSDMs) without impairing secretion of IL-1β, confirming previous findings that that glycine acts downstream of GSDMD pore formation (**Supplemental Figure 2B-D**).

**Figure 3.**
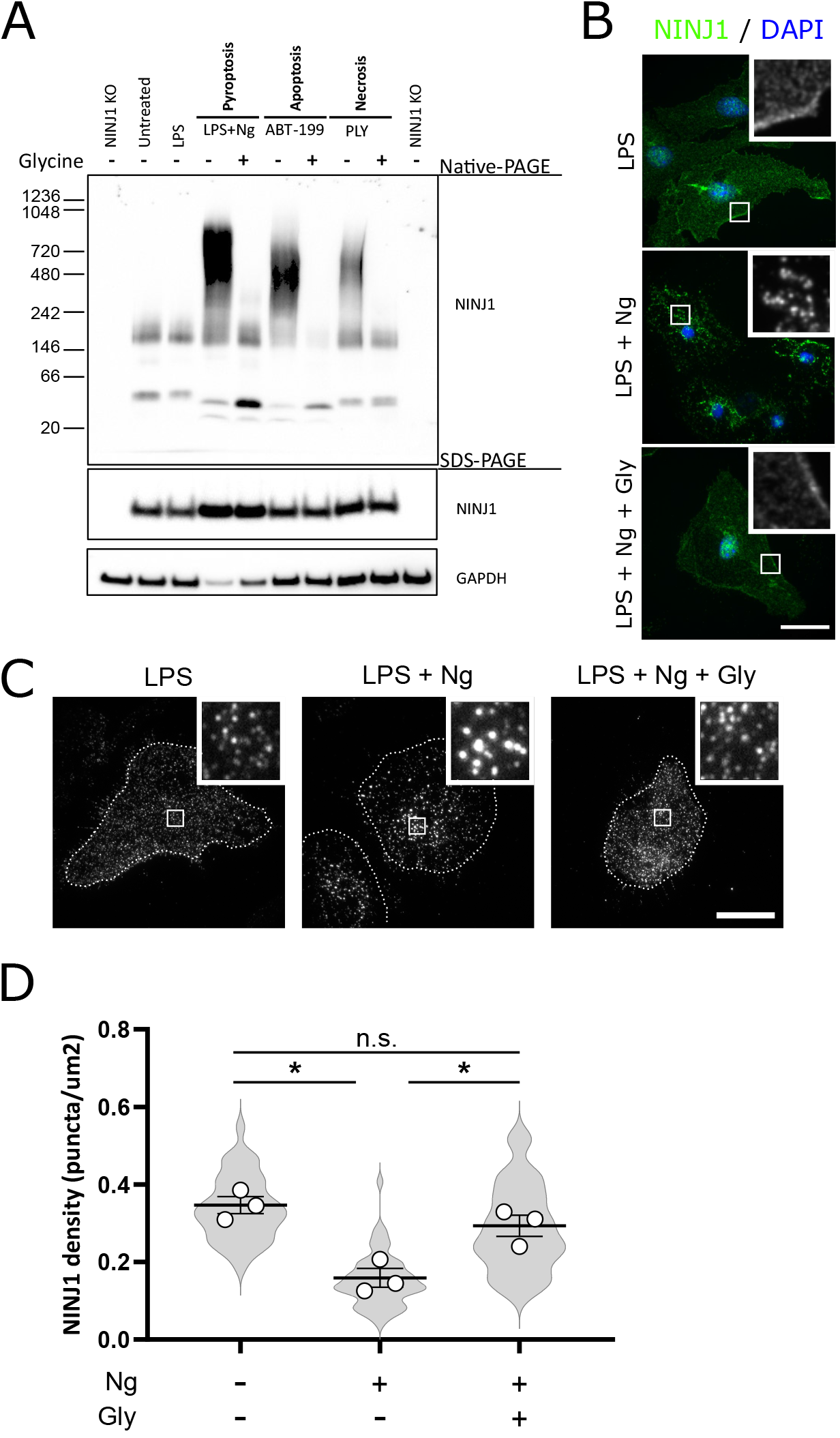
Glycine targets NINJ1 oligomerization to prevent membrane rupture in primary mouse macrophages. (**A**) Primary BMDM were induced to undergo pyroptosis (nigericin), apoptosis (venetoclax) or necrosis (pneumolysin) with or without glycine treatment. Native-PAGE analysis of endogenous NINJ1 demonstrates a shift to high molecular weight aggregate upon cell death stimulation, which is abrogated by glycine treatment. (**B**) Pyroptosis was induced in primary LPS-primed BMDM with nigericin with or without glycine. Immunofluorescence microscopy of native NINJ1 (green) reveals a redistribution of NINJ1 from diffuse plasma membrane staining to discrete puncta. NINJ1 does not redistribute in glycine-treated cells. Nuclei are labeled with DAPI (blue). Scale bar 15 μm. Inset shows magnified area demarcated by the white box. (**C**) Total internal reflection microscopy of NINJ1 in LPS-primed primary BMDM reveals that endogenous NINJ1 resides in discrete puncta within the plasma membrane. Cell membrane outline (dotted white line) was determined using fluorescently labeled cholera toxin subunit B (not shown) as a plasma membrane marker. Scale bar 20 μm. Inset shows magnified area demarcated by the white box. (**D**) Quantification of the density of NINJ1 puncta in LPS-primed primary macrophages at baseline or stimulated to undergo pyroptosis (nigericin 20 μM for 30 min) without or with glycine (5 mM). NINJ1 puncta become less dense upon pyroptosis induction, consistent with NINJ1 plasma membrane clustering. Glycine limits this redistribution. Violin plot of NINJ1 puncta density from 3 pooled independent experiments. Data points superimposed on the violin plots are the mean NINJ1 puncta densities for the 3 independent experiments (10-23 cells measured per replicate with >675 NINJ1 puncta identified per replicate). Bars represent mean ± SEM of the NINJ1 densities for the *n* = 3 independent experiments. * *P* < 0.05 by ANOVA with Tukey’s multiple comparison correction.

**Figure 4.**
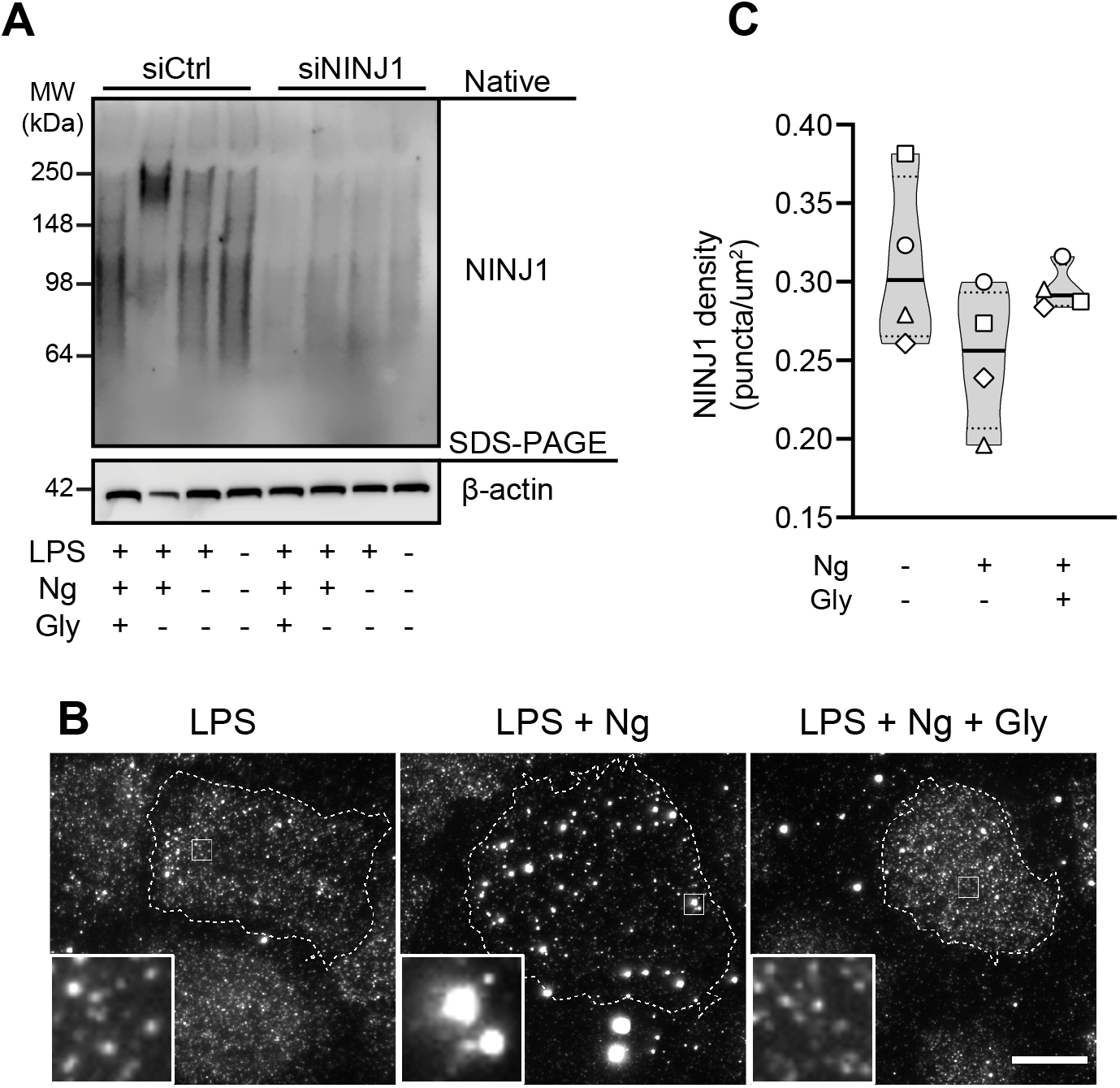
Glycine targets NINJ1 oligomerization to prevent membrane rupture in primary human macrophages. (**A**) Native-PAGE analysis of endogenous NINJ1 from hMDMs stimulated to undergo pyroptosis displays a shift to higher molecular weight, which is abrogated by glycine treatment. NINJ1 levels are almost absent in siNINJ1-treated cells (quantification in **Supplemental Figure 2A**) and no oligomerization is seen after LPS and nigericin treatment. (**B**) TIRF microscopy of LPS-primed primary hMDMs shows endogenous NINJ1 in discrete plasma membrane puncta. Cell membrane outline (dotted white line) was determined using fluorescently labeled cholera toxin subunit B (not shown) as a plasma membrane marker. Scale bar 20 μm. (**C**) Quantification of the NINJ1 puncta density in LPS-primed hMDMs at baseline or stimulated to undergo pyroptosis (nigericin 20.7 μM for 2 hours) without or with glycine (50 mM). Violin plot of NINJ1 puncta density quantified from images of cells from 4 independent donors. Mean of NINJ1 puncta densities from three cells per donor per condition shown as superimposed datapoints (different symbols are used for individual donors). Median and quartiles shown for each condition.

To corroborate these biochemical findings, we proceeded to evaluate native NINJ1 oligomerization by fluorescence spinning disk microscopy in primary BMDM induced to undergo pyroptosis. In unstimulated (not shown) and LPS-primed cells (**Figure 3B**), NINJ1 primarily localizes to the plasma membrane with some additional signal likely within the Golgi. Following induction of pyroptosis with nigericin in LPS-primed BMDM, NINJ1 redistributed into discrete puncta, which is consistent with clustering. Co-treatment with glycine prevented NINJ1 redistribution (**Figure 3B**).

Next, to confirm the plasma membrane localization of the observed puncta, we conducted total internal reflection (TIRF) microscopy of primary mouse BMDMs and hMDMs induced to undergo pyroptosis without or with glycine. TIRF microscopy allows for the preferential excitation of a thin (~150 nm) layer that includes the ventral plasma membrane of adherent cells, thereby eliminating background fluorescence from structures outside this focal plane (e.g. endomembrane compartments). In LPS-primed BMDMs and hMDMs, NINJ1 appears in discrete, small puncta that are diffusely distributed across the plasma membrane in both cell types (**Figure 3C**, **Figure 4B**). Upon induction of pyroptosis, the density of these puncta decreases (**Figure 3C-D**, **Figure 4B-C**), consistent with the clustering or aggregation of the basal dimers or trimers into high-order oligomers as seen on the native PAGE (**Figure 3A**, **Figure 4A**). Co-treatment with glycine abrogates the formation of these larger clusters (**Figure 3C-D**, **Figure 4B-C**).

### Discussion

The protective effect of glycine treatment is a pervasive feature of many cell death pathways, but with an unknown mechanism of action. Here, we position the transmembrane protein NINJ1 as a component of the elusive glycine target.

In our studies, we focused on fundamental cell death pathways in both mouse and human macrophages and show that glycine prevents NINJ1 clustering, a process required for plasma membrane rupture. Necroptosis represents another programmed cell death pathway, important to inflammatory responses, pathogen detection and tissue repair (Bertheloot et al., 2021). Unlike pyroptosis, necrosis and post-apoptosis lysis, NINJ1 does not appear to be needed for necroptosis-induced plasma membrane rupture (Kayagaki et al., 2021), potentially due to the membrane disrupting capacity of the necroptosis MLKL pore (Flores-Romero et al., 2020). This is particularly notable as glycine does not prevent TNF-induced necroptosis (Chen et al., 2014), further buttressing the association between NINJ1 and glycine.

Our data demonstrate that glycine treatment and genetic ablation or silencing of NINJ1 yield similar functional and morphological outcomes in both human and mouse macrophages subjected to lytic cell death programs. Mechanistically, our data suggest that glycine cytoprotection results from interference with NINJ1 clustering within the plasma membrane. This finding is further supported by the observation that glycine treatment of NINJ1 knockout cells confers no additional protection against cellular rupture. Future work should be directed at determining whether glycine directly or indirectly engages NINJ1. Indeed, how NINJ1 is activated by lytic cell death pathways, its clustering trigger, and method of plasma membrane disruption also remain important and outstanding questions in the field. Due to the ubiquity of lytic cell death pathways in human health and disease and the potency of glycine cytoprotection, answers to these questions will greatly advance the development of therapeutics against pathologic conditions associated with aberrant lytic cell death pathways.

## Methods

### Cells

Primary bone marrow derived macrophages (BMDM) were harvested from the femurs and tibia of wildtype mixed-sex cohorts of C57Bl/6 mice. The ends of cleaned bones were cut and centrifuged to collect bone marrow into sterile PBS. The cell suspension was washed with PBS and plated in DMEM with 10 ng mL^-1^ M-CSF (Peprotech Inc, 315-02). Following 5 days of culture, the BMDM were detached from the dishes with TBS with 5 mM EDTA, resuspended in fresh DMEM and plated. ASC-expressing RAW264.7 mouse macrophages (RAW-ASC; InvivoGen raw-asc) are engineered to stably express murine ASC, which is absent in the parental RAW 264.7 macrophage cell line (Pelegrin et al., 2008) and therefore are able to undergo pyroptosis. Immortalized BMDM expressing Cas9 were obtained from Dr Jonathan Kagan (Boston Children’s Hospital, Boston) and described previously (Evavold et al., 2018).

#### Isolation and differentiation of human primary macrophages

Buffy coats and pooled serum from healthy donors were provided by the blood bank at St. Olavs Hospital (Trondheim, Norway), after informed consent and with approval by the Regional Committee for Medical and Health Research Ethics (No. 2009/2245). Peripheral blood mononuclear cells (PBMCs) were isolated by density gradient using Lymphoprep™ (Axis-Shield) according to the manufacturer’s instructions. Monocytes were selected by α-CD14 bead isolation (Miltenyi Biotech) and seeded in RPMI 1640 supplemented with 10% serum and 10 ng/ml recombinant macrophage colony-stimulating factor (M-CSF; R&D Systems) for 3 days. At the third day, the medium was replaced with RPMI 1640 without M-CSF and siRNA treatment was initiated.

#### Induced pluripotent stem-cell (iPSC) derived macrophages (iPSDM) culture and production

iPSCs were obtained from European Bank for induced pluripotent Stem Cells (EBiSC,https://ebisc.org/about/bank), distributed by the European Cell Culture Collection of Public Health England (Department of Health, UK). iPSCs were maintained in Vitronectin XF (StemCell Technologies) coated plates with mTeSR™ Plus medium (StemCell Technologies). Cells were passaged with ReLeSR™ (StemCell Technologies) and plated in media containing 10 μM Rho-kinase inhibitor Y-27632 (StemCell Technologies). Embryonic body (EB) formation and myeloid differentiation is based on and adapted from a previously reported protocol (van Wilgenburg et al., 2013). Briefly, iPSCs were washed with PBS before Versene (Gibco-Thermo Fisher) was added and cells incubated at 37 °C for 5 minutes to generate a single cell suspension of iPSCs. Cells were counted, washed with PBS, and resuspended to a final concentration of 10 000 cells/50 μL in EB media: mTeSR™ Plus medium (StemCell Technologies), 50 ng/mL BMP-4 (R&D, 315-BP-010), 20 ng/mL SCF (R&D, 255-SC-050), 50 ng/mL VEGF (R&D, 293-VE-050). Cell suspension, supplemented with 10 μM Y-27632, was added into a 384-well Spheroid Microplate (Corning. #3830), 50 ul/well. The microplate was centrifuged at 1000 RPM for 3 minutes, and incubated at 37 °C, 5%CO_2_ for 4 days. EBs were fed at day 2 by adding 50 μL of fresh EB medium. After 4 days, EBs were harvested and seeded into a gelatin coated T175 flask in X-VIVO-15 (Lonza) media, supplemented with 100 ng/mL M-CSF (Prepotech, 300-25), 25 ng/mL IL-3 (Prepotech, 200-03), 2 mM glutamax (Gibco-Thermo Fisher), and 0.055 mM β-mercaptoethanol (Gibco-Thermo Fisher). Media was changed every week and EB-culture was renewed after 4 months. Monocytes produced were routinely checked with regards to phenotype using a defined flow cytometry panel of expected surface markers. Monocytes were harvested once a week and differentiated into macrophages in RPMI 1640 with 10% fetal calf serum and 100 ng/mL M-CSF for 5-7 days.

### NINJ1 knock-out cell line generation and siRNA silencing of NINJ1

RAW-ASC and iBMDM cells were transfected with custom CRISPR gRNA plasmid DNA (U6-gRNA:CMV-Cas-9-2A-tGFP) (Sigma-Aldrich) using FuGENE HD transfection reagent (Promega). The NINJ1 target region sequence was GCCAACAAGAAGAGCGCTG. Twenty-four hours later the cells were FACS sorted for GFP into 96 well plates. Individual colonies were expanded and tested for NINJ1 expression by western blot. NINJ1 knockdown in hMDMs was performed by siRNA using Lipofectamin RNAiMAX (Invitrogen) according to the manufacturer’s instructions. Cells were transfected with 20 nM pooled siRNA against NINJ1 (Thermo Scientific, HSS107188, HSS107190, HSS181529) or non-silencing control siRNA (Qiagen, 1027310). hMDMs were siRNA-treated two times (day 3 and day 5), before medium was changed to RPMI with 1% serum (day 6) and cells were allowed to rest overnight prior to stimulation. Knockdown of NINJ1 was confirmed by western blotting.

### Pyroptosis, apoptosis, and necrosis induction

Pyroptosis was activated in BMDM primed with (0.5 μg/ml) LPS from *E. coli* serotype 055:B5, which was first reconstituted at a stock concentration of 1 mg/ml. In primary BMDM, after 4.5 h of LPS priming, pyroptosis was induced with 20 μM nigericin (Sigma N7143; stock 10 mM in ethanol) for 30 min. Immortalized BMDM were primed with LPS for 3 h, followed by pyroptosis induction with 10 μM nigericin for 2 h. Apoptosis was induced in non-primed macrophages using the Bcl-2 inhibitor venetoclax (ABT-199, Tocris 6960) at 25 μM for 16 h. Necrosis was induced by treating non-primed macrophages with pneumolysin (0.5 μg/ml) for 15 min in iBMDM or 45 min in primary BMDM. Pneumolysin was obtained from Dr John Brumell (Hospital for Sick Children, Toronto). In hMDMs, pyroptosis was induced by priming with 1 μg/ml LPS from *E.coli* serotype 0111:B4 for 2 hours, followed by stimulation with 20.7 μM nigericin (Invivogen tlrl-nig; stock 5 mg/ml in ethanol) for 2 hours and 15 minutes. iPSDMs were stimulated with 200 ng/ml LPS for 3 hours and 20.7 μM nigericin for 1-1.5 hour.

Where indicated, mouse cells were treated with 5 mM glycine (Sigma, G7126) at the time of cell death induction. Human cells were pretreated with glycine (Millipore, 104201) at a concentration of 50 mM for at least 10 minutes before stimuli was added.

### LDH assay

BMDM were seeded at 200 000 cells per well in 12-well plates, treated as indicated, and cytotoxicity assayed by LDH release assay the following day. At the end of the incubations, cell culture supernatants were collected, cleared of debris by centrifugation for 5 min at 500 x *g*. The cells were washed once with PBS then lysed in lysis buffer provided in the LDH assay kit (Invitrogen C20300). Supernatants and lysates were assayed for LDH using an LDH colorimetric assay kit as per the manufacturer’s instructions (Invitrogen C20300). hMDMs were seeded at 500 000 cells per well in 12-well plates and differentiated as described above. iPSDMs were seeded at 180 000-450 000 cells per well in 12-well plates and differentiated as described above. After cells were stimulated as described in figure legends, supernatants were harvested and LDH release was assayed (Invitrogen C20301). Briefly, equal amounts of supernatant and reaction mixture was mixed and incubated for 30 minutes, followed by absorbance reads at 490 and 655 nm. To calculate the LDH release, the 655 nm background was subtracted from the 490 nm absorbance reads before further processing.

### Fluorescence microscopy

<u>Mouse macrophages</u> were cultured on 18 mm glass coverslips in 12-well plates at 150 000 cells per well. Cell were treated as indicated in the figure legends, washed with PBS, fixed in 4% paraformaldehyde in PBS at room temperature for 15 min, and permeabilized with 0.1% Tween-20. Cells were then blocked in PBS supplemented with 10% donkey serum and 0.1% Tween-20 for 1 h at room temperature prior to overnight incubation with primary antibody at 4 °C. Rabbit monoclonal anti-mouse NINJ1 was used at 10 μg/ml. Next, cells were washed three times with PBS supplemented with 10% donkey serum prior to the addition of secondary antibody at room temperature for 1 h. Nuclei were labeled using DAPI containing mounting medium (ProLong Diamond Antifade Mountant, Invitrogen P36961). Cells were imaged by spinning disk confocal microscopy (Quorum) on a Zeiss Axiovert 200M microscope with a 63x objective and an additional 1.5x magnifying lens. Images were acquired by a CCD camera (Hamamatsu Photonics) using Volocity acquisition software. For total internal reflection microscopy, cells were co-stained with recombinant Cholera toxin subunit B conjugated to Fluor488 (Invitrogen C34775) at 1 μg/mL for less than 1 minute prior to fixing and staining for NINJ1 as described above. The Cholera toxin subunit B served as a membrane marker to optimize plasma membrane visualization. TIRF imaging of mouse macrophages was conducted on a Zeiss Axio Observer Z1 microscope with a 100x objective and Andor iXon3 885 detector. Rabbit monoclonal anti-mouse NINJ1 were obtained from Drs Nobuhiko Kayagaki and Vishva Dixit (Genentech Inc, San Francisco). Where indicated, images of live cell morphology were captured using the epifluorescence mode and with FM™ 4-43 (Invitrogen, T3166) added right before imaging.

<u>Human primary macrophages</u> were differentiated at 30 000 cells per well in glass-bottom 96-well plates (Cellvis, P96-1.5H-N). After stimuli, the cells were washed with PBS and thereafter stained with Cholera toxin subunit B Alexa Fluor 488 conjugate at 1 μg/ml in PBS for less than 1 minute. Cells were then fixed in 4% PFA in PBS at room temperature for 15 minutes, followed by washing three times with PBS. We permeabilized the cells with PBS containing 0,01% saponin for 10 minutes at room temperature, and thereafter blocked with PBS/0,01% saponin/20% pooled human serum for 3 hours. Cells were incubated with human NINJ1 antibody (R&D Systems, MAB5105) at 10 μg/ml in PBS/0,01% saponin/1% pooled human serum at 4 °C for 41 hours, and then washed three times with PBS/0,01% saponin/1% pooled human serum. Following this, we incubated the cells with far-red fluorescent rabbit anti-mouse antibody (Invitrogen, A27029) at 2 μg/ml in PBS/0,01% saponin/1% pooled human serum at room temperature for 1 hour. Cells were washed three times with PBS/0,01% saponin/1% pooled human serum and thereafter three times with PBS before imaging. TIRF imaging of human macrophages was done on a Zeiss TIRF 3 microscope with a 63x objective and a Hamamatsu ORCA Fusion sCMOS detector. Images of morphology in live cells were captured using the epifluorescence mode and with FM™ 1-43 (Invitrogen, T3163) added at 5 μg/ml right before imaging.

### Native and SDS-PAGE

Mouse macrophages were lysed with native-PAGE lysis buffer (150 mM NaCl, 1% Digitonin, 50 mM Tris pH7.5, and 1 x Complete Protease Inhibitor). Following centrifugation at 20 800 x *g* for 30 min, lysates were mixed with 4X NativePAGE sample buffer and Coomassie G-250 (ThermoFisher) and resolved using NativePAGE 3-12% gels. For SDS-PAGE, cells were washed with 1x PBS and lysed in RIPA lysis buffer containing protease inhibitors (Protease inhibitor tablet, Pierce A32955). Proteins were resolved using NuPAGE 4-12% Bis-Tris gels (Invitrogen), transferred to PVDF membranes and immunoblotted. GAPDH (Santa Cruz, sc-25778; 1:1000) was used for a loading control.

hMDMs and iPSDMs were lysed with 1X NativePAGE™ Sample Buffer containing 1% Digitonin. After centrifugation at 21 000 x g for 30 min, protein concentration of lysates was measured using Pierce™ BCA Protein Assay Kit (ThermoFisher), to ensure equal loading of proteins. For Native-PAGE on hMDMs, lysates were mixed with Novex™ Tris-Glycine Native Sample Buffer (Invitrogen) and resolved using Novex™ 4-12% Tris-Glycine gels (Invitrogen). For Native-PAGE on human iPSDMs, lysates were mixed with Coomassie G-250 (ThermoFisher) and resolved using NativePAGE 3-12% Bis-Tris gels (Invitrogen). For SDS-PAGE, lysates were mixed with DTT and NuPAGE LDS sample buffer (Invitrogen), denatured for 5 min at 95 °C, and resolved using NuPAGE 4-12% Bis-Tris gels (Invitrogen). Proteins were transferred to PVDF membranes and immunoblotted. NINJ1 (R&D Systems, MAB5105) was used to determine NINJ1 expression, while β-actin (Cell Signaling Technology, 8457) was used as loading control.

### Cytokine measurements

Supernatants from human iPSDMs were assessed for IL-1β (R&D Systems, DY201) by ELISA (R&D Systems) according to the protocol of the manufacturer.

### Quantification and Statistics

TIRF microscopy images of mouse macrophages were analyzed in Imaris (Oxford Instruments) software. The Spot tool was used to identify NINJ1 puncta with parameters (estimated puncta *XY* diameter, region type, quality filter) set using images from the LPS-primed group and applied to all samples across all experiments. Cells were contoured using the surfaces tool to measure cell area. The density of NINJ1 puncta was calculated on a per-cell basis by dividing the total number of puncta within the contoured plasma membrane area. TIRF microscopy images of human primary macrophages were analyzed in FIJI, applying thresholds (triangle algorithm) and using the analyze particles tool to segment cells in the Cholera toxin subunit B channel. The find maxima command was used to identify NINJ1 puncta within these cell outlines, and the measurement tool to measure area of the cell outlines. The density of NINJ1 puncta was calculated on a per-cell basis by dividing the number of puncta within a cell outline with its area.

Statistical testing was calculated using Prism 8.3.1 or 9.0 (GraphPad Software Inc, La Jolla, USA). Experiments with more than two groups were tested using ANOVA with Tukey’s multiple comparison test. Unless otherwise indicated, presented data are representative of at least three independent experiments or donors and are provided as mean ± SEM.

### Animal studies

All animal studies were approved by the Hospital for Sick Children Animal Care Committee.

### Human studies

All human studies were conducted according to the principles expressed in the Helsinki Declaration and informed consent was obtained from all subjects prior to sample collection.

## Author contributions

JPB and RSRS conducted experiments, acquired and analyzed data, and assisted in writing the manuscript. They contributed equally and are listed alphabetically. AV, MB, and BK conducted experiments, acquired and analyzed data, and assisted in writing the manuscript. NMG and THF designed research studies, analyzed data, and assisted in writing the manuscript. BES supervised the project, designed research studies, analyzed data, and wrote the manuscript.

## Acknowledgements

This work was supported by a Mentored Research Award from the *International Anesthesia Research Society* and an Early Investigator Award from the Department of Anesthesiology & Pain Medicine, University of Toronto to BES, and by grants (287696, 223255) from the Research Council of Norway to THF. We thank Dr Nobuhiko Kayagaki and Dr Vishva Dixit for providing the rabbit monoclonal anti-mouse NINJ1 antibody, Dr Jonathan Kagan for providing the immortalized bone marrow derived macrophages expressing Cas9, Dr John Brumell for providing pneumolysin, and Anne Marstad for technical assistance. All imaging of the human cells was performed at the Cellular and Molecular Imaging Core Facility at NTNU. Graphical abstract created with BioRender.com. The authors declare no competing financial interests.

---

**Supplemental Figure 1.**
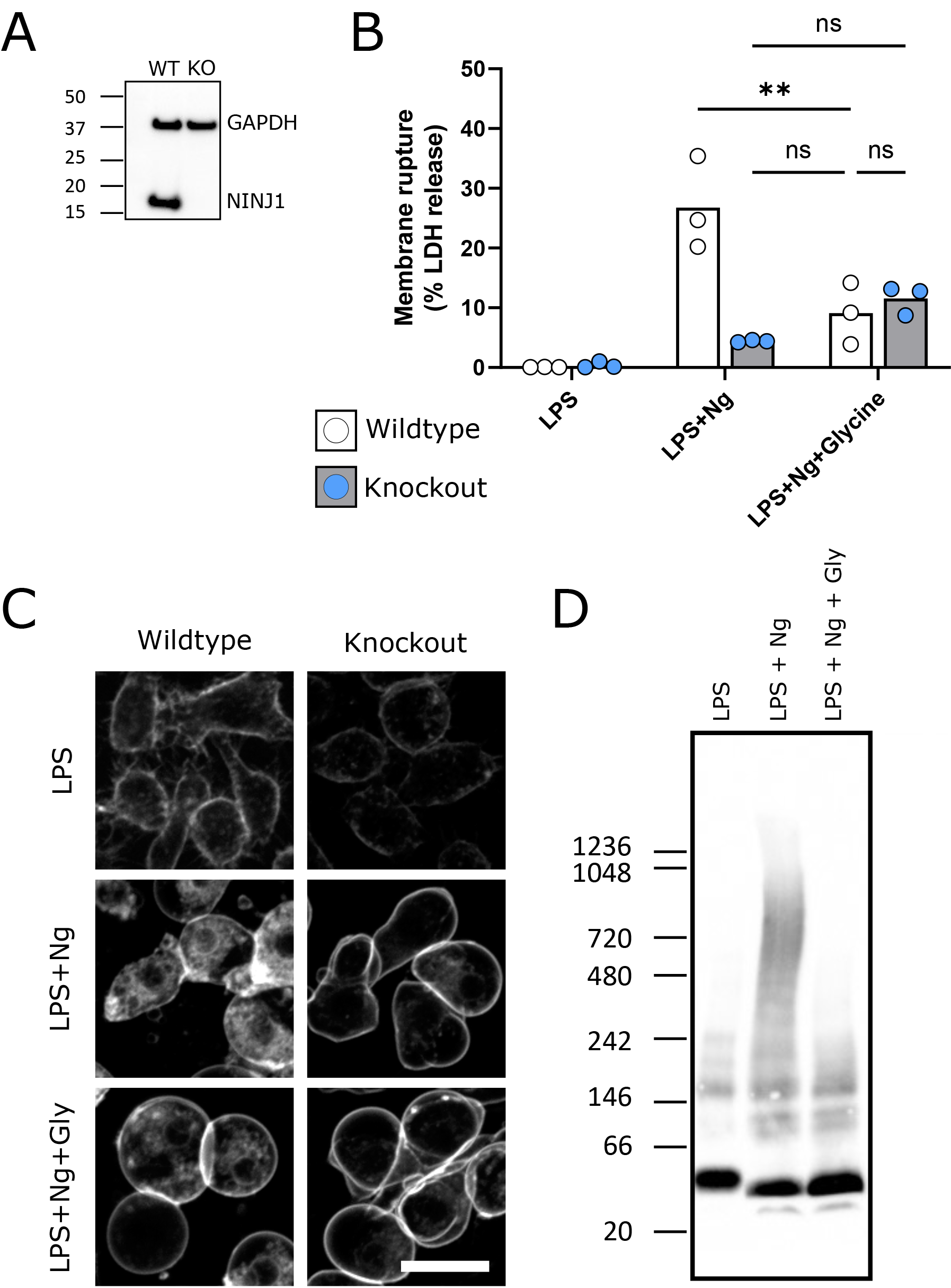
Glycine cytoprotection phenocopies NINJ1 knockout and prevents NINJ1 aggregation in a mouse macrophage cell line. Using a CRISPR-Cas9 system, NINJ1 was knocked out of a RAW mouse macrophage cell line engineered to constitutively express ASC (RAW-ASC) and therefore have the capacity to undergo pyroptosis. (**A**) Western blot demonstrating NINJ1 knockout compared with the parental line. GAPDH is shown as a loading control. (**B**) Wildtype and NINJ1 knockout were induced to undergo pyroptosis with LPS + nigericin treatment with or without glycine. Supernatant LDH was evaluated as a marker of cytotoxicity by colorimetric assay. Cytotoxicity was decreased in glycine-treated wildtype cells comparably to NINJ1 knockout. Glycine treatment of NINJ1 KO cells provided no additional protection to knockout cells without glycine. * P < 0.05 by two-sided ANOVA with Tukey’s multiple comparison correction. (**C**) LPS-primed wildtype RAW-ASC induced to undergo pyroptosis in the presence of glycine demonstrate similar plasma membrane ballooning to NINJ1 knockout RAW-ASC cells induced to undergo pyroptosis. Membrane ballooning is shown in live cells labelled with the plasma membrane dye FM1-43. Scale bar 20 μm. (**D**) Native-PAGE analysis of endogenous NINJ1 in RAW-ASC macrophages demonstrates a shift to high molecular weight aggregate upon pyroptosis stimulation, which is abrogated by glycine treatment.

**Supplemental Figure 2.**
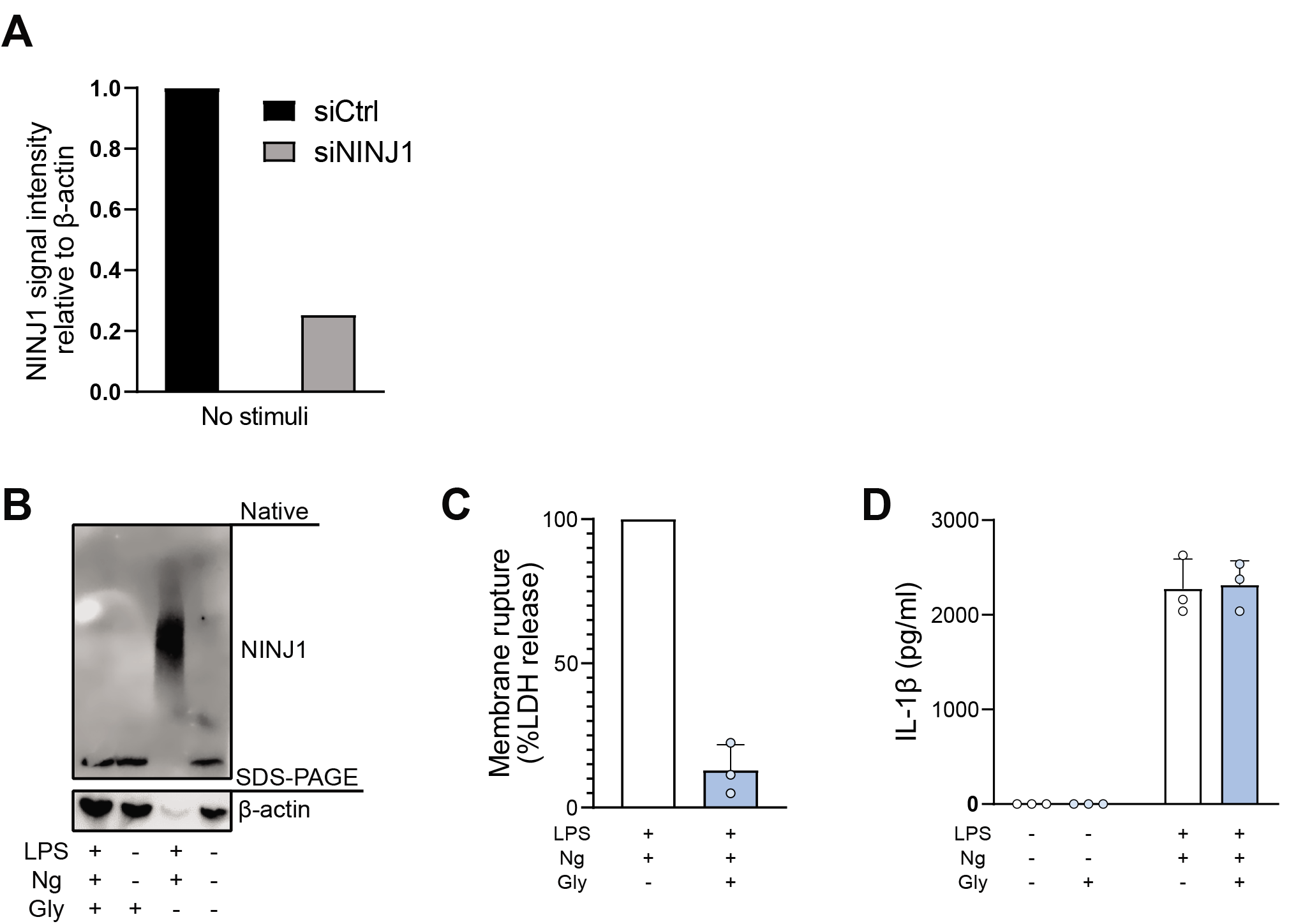
Glycine prevents NINJ1 aggregation and pyroptotic lysis in induced pluripotent stem-cell derived human macrophages without affecting IL-1β secretion. (**A**) Quantification of NINJ1 native-PAGE signal from unstimulated hMDM lysates confirm NINJ1 silencing in siNINJ1-treated cells compared to siCtrl-treated cells. (**B-D**) Induced pluripotent stem-cell (iPSC) derived macrophages (iPSDMs) were LPS-primed and stimulated to undergo pyroptosis with nigericin in the presence or absence of 50 mM glycine. (**B**) Native-PAGE analysis of endogenous NINJ1 in iPSDMs display a shift to higher molecular weight when cells are stimulated to undergo pyroptosis, which is abrogated by glycine pretreatment. (**C**) LDH release was measured in the supernatants to assess cell rupture and was reduced with glycine co-treatment. Data are expressed as % of LDH in supernatant from LPS and nigericin-treated cells in *n* = 3 independent experiments. Data points from each experiment is shown along with their mean and standard deviation. (**D**) IL-1β release from pyroptotic cells was not altered by the presence of glycine. Data points from three technical replicates from one experiment shown along with their mean and standard deviation.

